# Fast and precise single-cell data analysis using hierarchical autoencoder

**DOI:** 10.1101/799817

**Authors:** Duc Tran, Hung Nguyen, Bang Tran, Carlo La Vecchia, Hung N. Luu, Tin Nguyen

**Affiliations:** Department of Computer Science and Engineering, University of Nevada Reno, Reno, NV, USA; Department of Clinical Sciences and Community Health, University of Milan, Milan, Italy; Division of Cancer Control and Population Sciences, Hillman Cancer Canter, University of Pittsburgh Medical Center, Pittsburgh, PA, USA; Department of Epidemiology, University of Pittsburgh Graduate School of Public Health, Pittsburgh, PA, USA

## Abstract

A primary challenge in single-cell RNA sequencing (scRNA-seq) studies comes from the massive amount of data and the excess noise level. To address this challenge, we introduce a hierarchical autoencoder that reliably extracts representative information of each cell. In an extensive analysis, we demonstrate that the approach vastly outperforms state-of-the-art techniques in many research sub-fields of scRNA-seq analysis, including cell segregation through unsupervised learning, visualization of transcriptome landscape, cell classification, and pseudo-time inference.

---

Advances in microfluidics and sequencing technologies have allowed us to monitor biological systems at single-cell resolution.^1, 2^ This comprehensive decomposition of complex tissues holds enormous potential in developmental biology and clinical research.^3–5^ However, the ever-increasing number of cells, technical noise, and high dropout rate pose significant computational challenges in scRNA-seq analysis.^6–8^ These challenges affect both analysis accuracy and scalability, and greatly hinder our capability to extract the wealth of information available in single-cell data.

To detach noise from informative biological signals, we have developed a new analysis framework, called single-cell Decomposition using Hierarchical Autoencoder (scDHA), that consists of two core modules (Figure 1a). The first module is a non-negative kernel autoencoder that provides a non-negative, part-based representation of the data. Based on the weight distribution of the encoder, scDHA removes genes or components that have insignificant contribution to the representation. The second module is a Stacked Bayesian Self-learning Network that is built upon the Variational Autoencoder^9^ to project the data onto a low dimensional space (see Online Methods). Using this informative and compact representation, many analyses can be performed with high accuracy and tractable time complexity (mostly linear or lower complexity).

**Figure 1.**
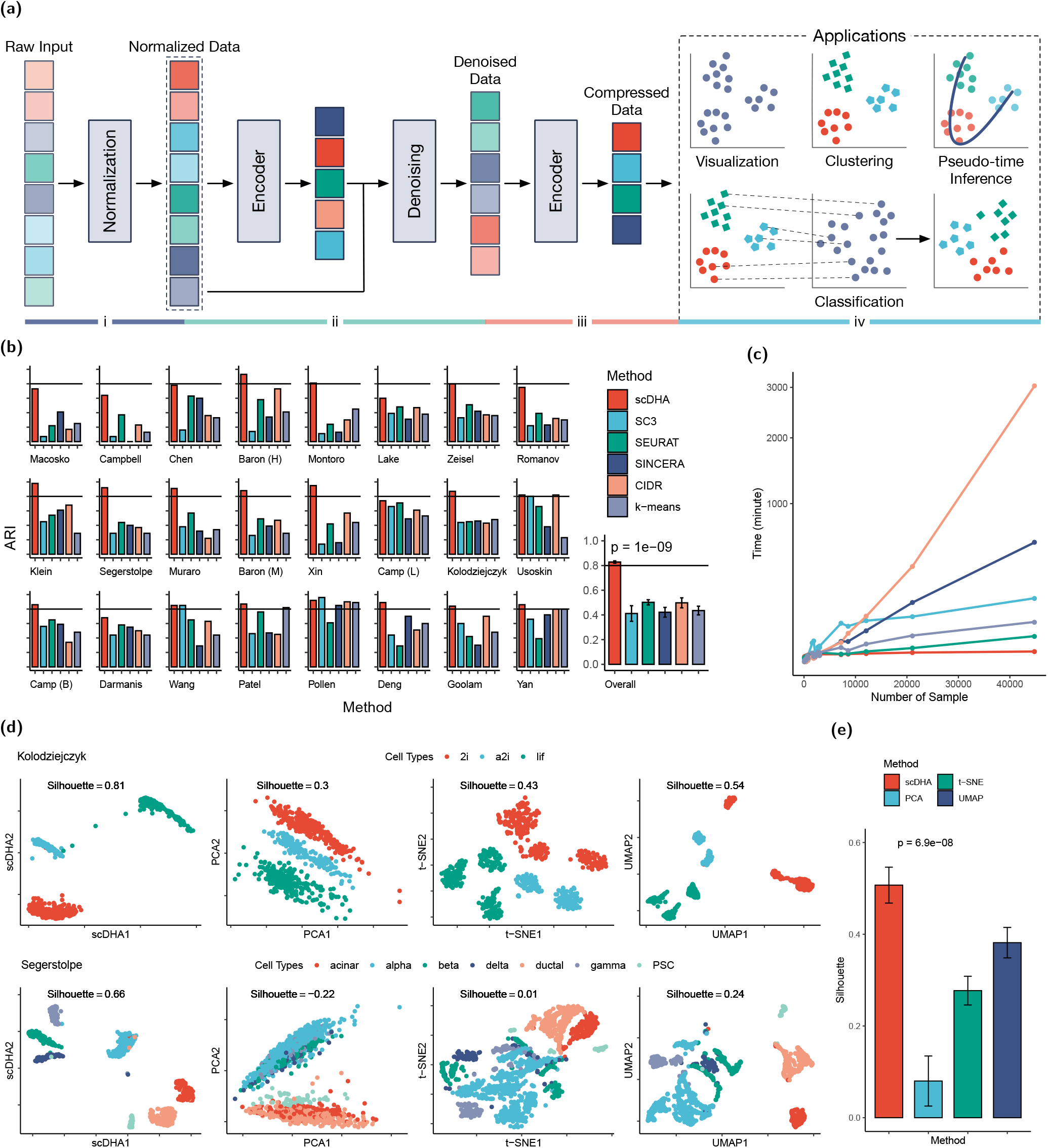
Overview of scDHA architecture and analysis performance on 24 scRNA-seq datasets. (a) Schematic overview of scDHA and applications: cell segregation through unsupervised learning, visualization, pseudo-temporal ordering, and cell classification. (b) Clustering performance of scDHA, SC3, SEURAT, SINCERA, CIDR, and k-means measured by adjusted Rand index (ARI). The first 24 panels show the ARI values obtained for individual datasets while the last panel shows the average ARIs and their variance (vertical segments). (c) Running time of the clustering methods, each using 10 cores. scDHA is the fastest among the six methods. (d) Color-coded representation of the Kolodziejczyk and Segerstolpe datasets using scDHA, PCA, t-SNE, and UMAP (from left to right). For each representation, we report the silhouette index, which measures the cohesion among the cells of the same type, as well as the separation between different cell types. (e) Average silhouette values (bar plot) and their variance (vertical lines). scDHA significantly outperforms other dimension reduction methods by having the highest silhouette values (*p* = 6.9 × 10^−8^ using Kruskal-Wallis test).

In one joint framework, the scDHA software package conducts cell segregation through unsupervised learning, dimension reduction and visualization, cell classification, and time-trajectory inference. We will show that scDHA outperforms state-of-the-art methods in all four sub-fields.

## Cell segregation

Defining cell types through unsupervised learning is considered the most powerful application of scRNA-seq.^7^ This has led to the creation of a number of atlas projects,^10, 11^ which aim to build the references of all cell types in model organisms at various developmental stages. We assess the performance of scDHA in clustering using 24 scRNA-seq datasets with known cell types (see Online Method for details of each dataset). The true class information of these datasets is only used *a posteriori* to assess the results. We compare scDHA with four methods that are widely used for single-cell clustering: SC3,^12^ SEURAT,^13^ SINCERA,^14^ and CIDR.^15^ We also include k-mean as the reference method.

Since the true cell types are known in these datasets, we use adjusted Rand index (ARI)^16^ to assess the performance of the six clustering methods. Figure 1b shows the ARI values obtained for each dataset, as well as the average ARI and their variance. scDHA outperforms all other methods by not only having the highest average ARI, but also being the most consistent method. The average ARI of scDHA across all 24 datasets is 0.83 with very low variability. The second best method, CIDR, has an average ARI of only 0.5. Kruskal-Wallis test also indicates that the ARI values of scDHA are significantly higher than the rest with a p-value of 10^−9^.

To perform a more comprehensive analysis, we calculate the normalized mutual information (NMI) and Jaccard index (JI) for each method (Supplementary Section 1 and Tables 1–3). Regardless of the assessment metrics, scDHA consistently outperforms all other methods. At the same time, scDHA is also the fastest among the six methods (Figure 1c and Supplementary Table 4) with an average running time of two minutes per analysis. For the Macosko dataset with 44 thousand cells, scDHA finishes the analysis in less than five minutes. On the contrary, it takes CIDR more than two days (3000 minutes) to finish the analysis of this dataset. In summary, scDHA outperforms other clustering methods in terms of both accuracy and scalability.

## Dimension reduction and visualization

Dimension reduction techniques aim at representing high-dimensional data in a low-dimensional space while preserving relevant structure of the data. Non-linear methods,^17^ including Isomap,^18^ Diffusion Map,^19^ t-SNE,^20^ and UMAP,^21^ have been recognized as efficient techniques to avoid overcrowding due to the large number of cells, while preserving the local data structure. Among these, t-SNE is the most commonly used technique while UMAP is a recent method. Here we demonstrate that scDHA is more efficient than both t-SNE and UMAP, as well as the classical principal component analysis (PCA) in visualizing single-cell data. We test the four techniques on the same 24 single-cell datasets described above. Again, cell type information is not given as input to any algorithm.

The top row of Figure 1d shows the color-coded representations of the Kolodziejczyk dataset, which consists of three mouse embryo stem cells: *2i*, *a2i*, and *lif*. The classical PCA simply rotates the orthogonal coordinates to place dissimilar data points far apart in the two-dimensional (2D) space. In contrast, t-SNE focuses on representing similar cells together in order to preserve the local structure. In this analysis, t-SNE mistakenly splits each of the two classes *2i* and *a2i* into two smaller groups, and *lif* class into three groups. The transcriptome landscape represented by UMAP is similar to that of t-SNE, in which UMAP also mistakenly splits cells of the same types into smaller groups. In contrast, scDHA provides a clear representation of the data, in which cells of the same type are grouped together and cells of different types are well-separated.

The lower row of Figure 1d shows the visualization of the Sergerstolpe dataset (human pancreas). The landscape of UMAP and t-SNE are better than that of PCA. In both representations, the cell types are separable. However, the cells are overcrowded and many cells from different classes overlap. In addition, the *alpha* and *gamma* cells are mistakenly split into smaller groups. For this dataset, scDHA again better represents the data by clearly showing the transcriptome landscape with separable cell types.

To quantify the performance of each method, we calculate the silhouette index (SI)^22^ of each representation using true cell labels. This metric measures the cohesion among the cells of the same type and the separation among different cell types. For both datasets shown in Figure 1d, the SI values of scDHA are much higher than those obtained for PCA, t-SNE, and UMAP. The visualization and SI values of the 22 other datasets are shown in Supplementary Figures 1–6 and Table 5. The average SI values obtained across the 24 datasets are shown in Figure 1e. Overall, scDHA consistently and significantly outperforms other methods (*p* = 6.9 × 10^−8^).

## Cell classification

*De novo* identification of cell types and building comprehensive atlases are a problem of unsupervised learning. Once the cellular subpopulations have been determined and validated, classification techniques can be used to determine the composition of new datasets by classifying cells into discrete types. We assess scDHA’s classification capability by comparing it with four methods that are dominant in machine learning: XGBoost,^23^ Random Forest (RF),^24^ Deep Learning (DL),^25^ and Gradient Boosting Machine (GBM).^26^

We test these methods using five datasets: Baron (8,569 cells), Segerstolpe (2,209 cells), Muraro (2,126 cells), Xin (1,600 cells), and Wang (457 cells). All five datasets are related to human pancreas and thus have similar cell types. In each analysis scenario, we use one dataset as training and then classify the cells in the remaining four datasets. For example, we first train the model on Baron and then test it on Segerstolpe, Muraro, Xin, and Wang. Next, we train the model on Segerstolpe and test on the rest, etc. The accuracy of each method is shown in Figure 2 and Supplementary Table 7.

**Figure 2.**
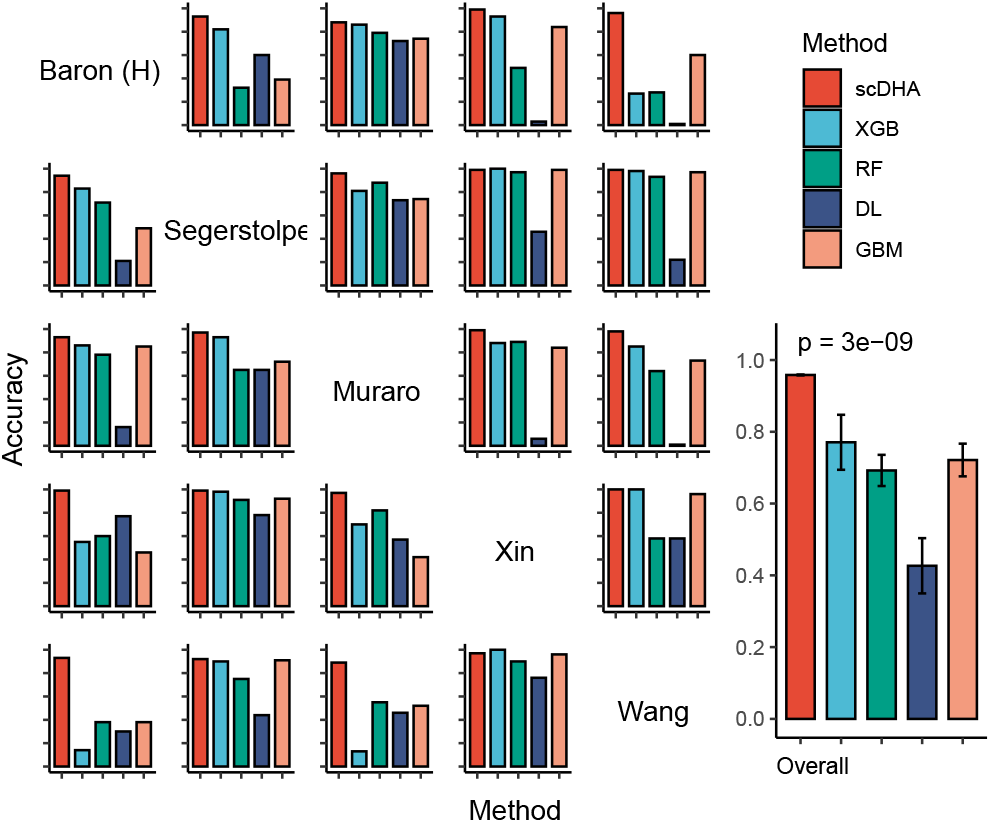
Classification accuracy of scDHA, XGBoost, Random Forest (RF), Deep Learning (DL), Gradient Boosted Machine (GBM) using 5 human pancreatic datasets. In each scenario (row), we use one dataset as training and the rest as testing, resulting in 20 train-predict pairs. The accuracy values of scDHA are significantly higher than those of other methods (*p* = 3 × 10^−9^ using Kruskal-Wallis).

Overall, scDHA is accurate across all 20 combinations with accuracy ranging from 0.88 to 1. Compared to other methods, scDHA is superior with an average accuracy of 0.96 for scDHA compared to 0.77, 0.69, 0.43, and 0.72 for XGB, RF, DL, and GBM, respectively. In addition, scDHA is very consistent, while the performance of existing methods fluctuates from one analysis to another, especially when the testing dataset is much larger than the training dataset. For example, when the testing set (Baron) is 20 times larger than the training set (Wang), the accuracy of existing methods is close to 30%, while scDHA achieves an accuracy of 0.93. A Kruskal-Wallis test also confirms that the accuracy values of scDHA are significantly higher than the rest (*p* = 3 × *p* = 10^−9^). Regarding time complexity, scDHA is the fastest with an average running time of two minutes per analysis (Supplementary Figure 7).

## Time-trajectory inference

Cellular processes, such as cell cycle, proliferation, differentiation, and activation,^27, 28^ can be modeled computationally using trajectory inference methods. These methods aim at ordering the cells along developmental trajectories. Among a number of trajectory inference tools, Monocle,^29^ TSCAN,^30^ and Slingshot^31^ are considered state-of-the-art and are widely used for pseudo-temporal ordering. Here we test scDHA and these methods using three mouse embryo development datasets: Yan, Goolam, and Deng. The true developmental stages of these datasets are only used *a posteriori* to assess the performance of the methods.

Figure 3a shows the Yan dataset in the first two t-SNE components. The smoothed lines shown in each panel indicate the time-trajectory of scDHA (left) and Monocle (right). The trajectory inferred by scDHA accurately follows the true developmental stages: it starts from zygote, going through 2cell, 4cell, 8cell, 16cell, and then stops at the blast class. On the contrary, the trajectory of Monacle goes directly from zygote to 8cell before coming back to 2cell. Figure 3b shows the cells ordered by pseudo-time. The time inferred by scDHA is strongly correlated with the true developmental stages. On the other hand, Monocle fails to differentiate between zygote, 2cell, and 4cell. To quantify how well the inferred trajectory explains the developmental stages, we also calculate the R-squared. scDHA outperforms Monocle by having a higher R-squared (0.93 compared to 0.84).

**Figure 3.**
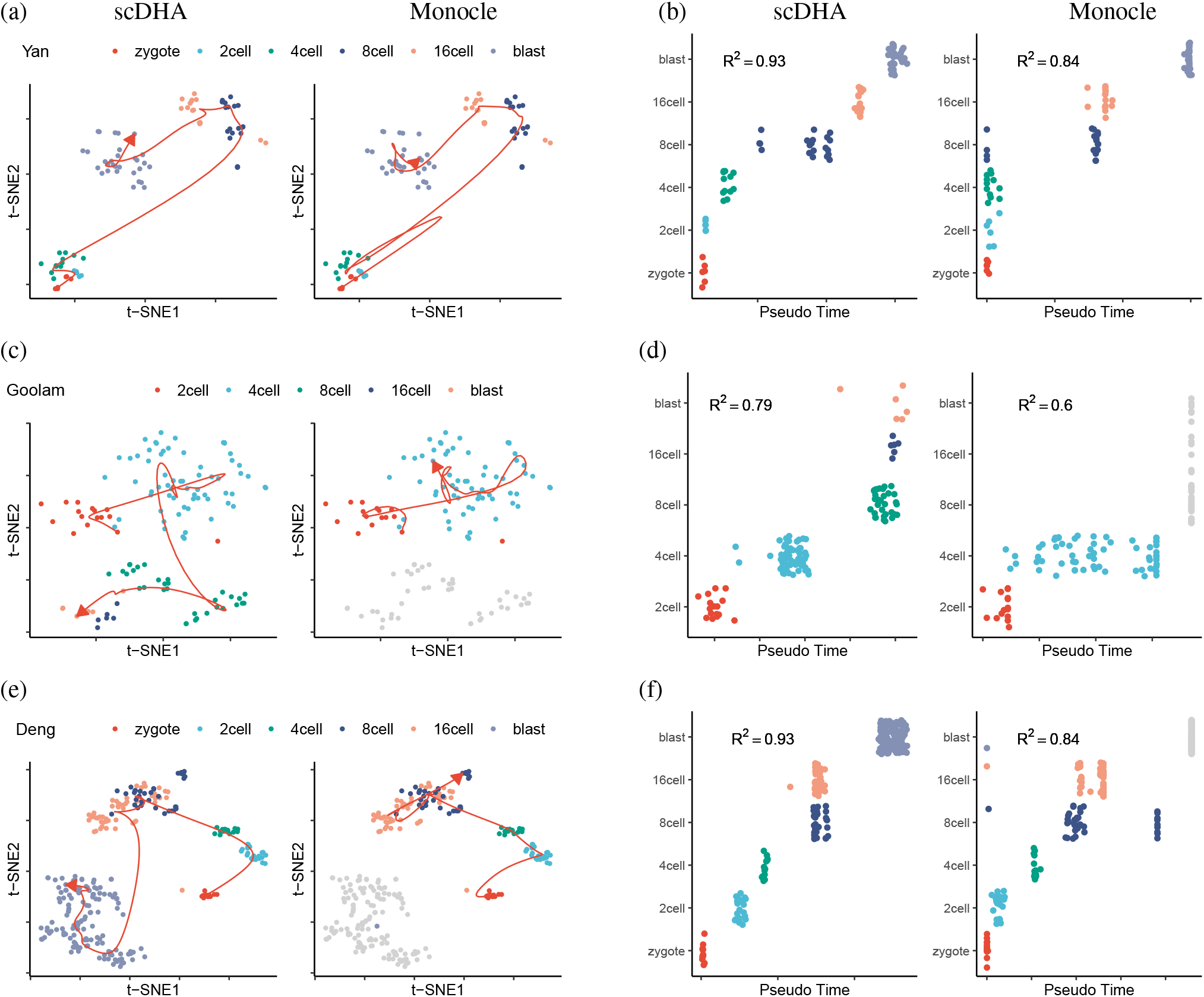
Pseudo-time inference of three mouse embryo development datasets (Yan, Goolam, and Deng) using scDHA and Monocle. (a) Visualized time-trajectory of the Yan dataset in the first two t-SNE dimensions using scDHA (left) and Monocle (right). (b) Pseudo-temporal ordering of the cells in the Yan dataset. The horizontal axis shows the inferred time for each cell while the vertical axis shows the true developmental stages. (c,d) Time-trajectory of the Goolam dataset. Monocle is unable to estimate the time for most cells in 8-cell, 16-cell, and blast (colored in gray). (e,f) Time-trajectory of the Deng dataset. Monocle is unable to estimate the pseudo time for most blast cells.

Figure 3c,d show the results of the Goolam dataset. scDHA correctly reconstructs the time-trajectory whereas Monocle fails to estimate pseudo-time for 8cell, 16cell, and blast cells (colored in gray). Monocle assigns an “infinity” value for these cell classes. Figure 3e,f show the results obtained for the Deng dataset. Similarly, the time-trajectory inferred by scDHA accurately follows the developmental stages whereas Monocle could not estimate the time for half of the cells. The results of TSCAN and Slingshot are shown in Supplementary Figures 8, 9). scDHA outperforms all three methods by having the highest R-squared values in every single analysis.

In summary, we have introduced a powerful framework for scRNA-seq data analysis. We have shown that the framework can be utilized for both upstream and downstream analyses, including *de novo* clustering of cells, visualizing the transcriptome landscape, classifying cells, and inferring pseudo-time. We demonstrate that scDHA outperforms state-of-the-art techniques in each research sub-field. Although we focus on single-cell as an example, scDHA is flexible enough to be adopted in a range of research areas, from cancer to obesity to aging to any other area that employs high-throughput data.

## Online Method

### Data and pre-processing

The 24 single-cell datasets used in our data analysis are described in Table 1. We download the Montoro dataset from Broad Institute Single Cell Portal (portals.broadinstitute.org/single_cell/study/SCP163/airway-epithelium), and the other 23 datasets from the website of Hemberg Group at the Sanger Institute (hemberg-lab.github.io/scRNA.seq.datasets). We removed samples with ambiguous label from these datasets. Specifically, we removed cells with label “zothers” from Chen, “Unknown” from Camp (Brain), “dropped” from Wang, and “not applicable” from Segerstolpe. The only processing step we did is to perform log transformation (base 2) to rescale the data if the range of the data is larger than 100.

**Table 1.**
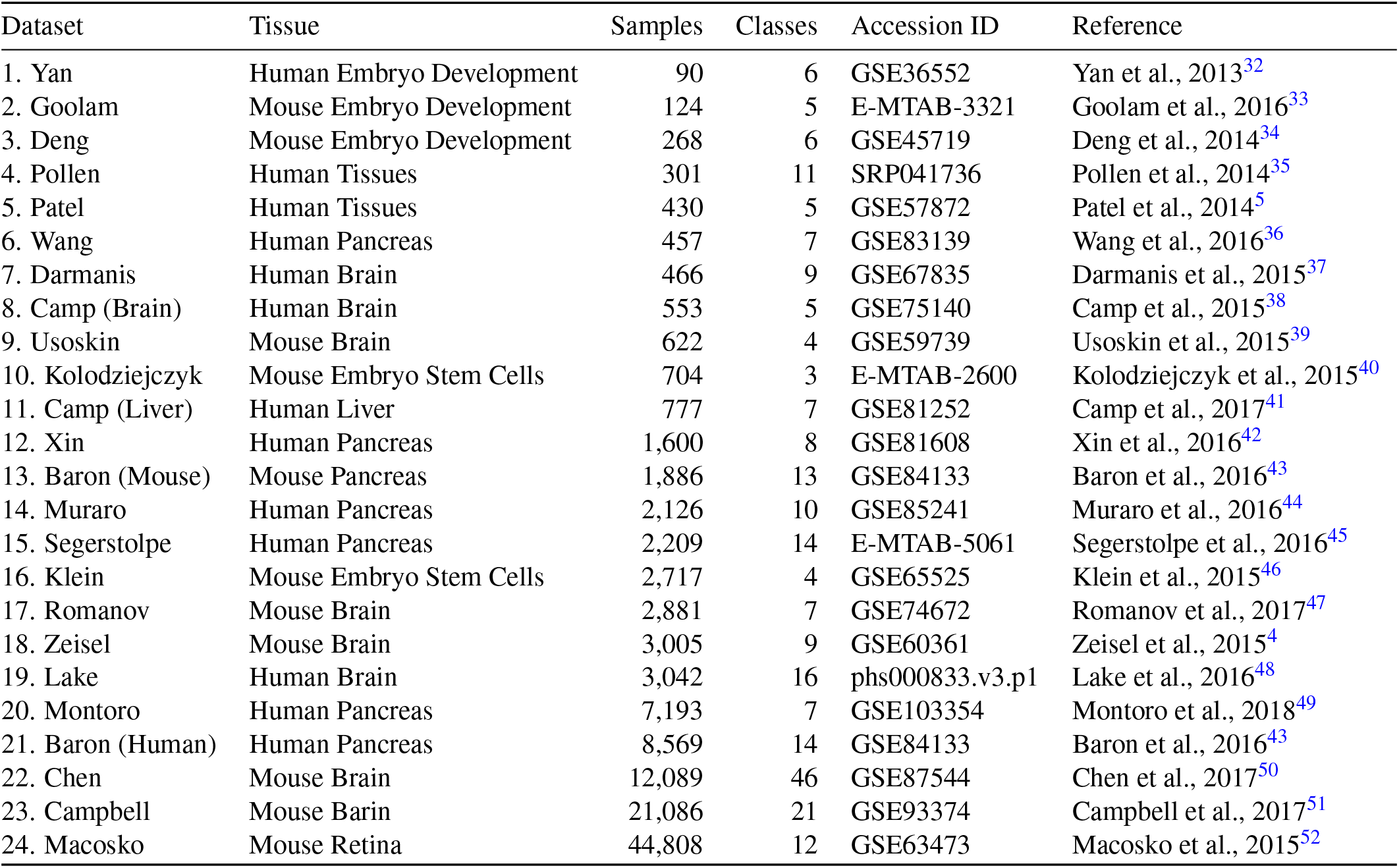
Description of the 24 single-cell datasets used to assess the performance of computational methods. The first two columns describe the name and tissue while the next three columns show the number of samples, number of cell types, and accession ID.

### Software package and setting

In our analysis, we followed the instruction and tutorial provided by the authors of each software package. We used the default parameters of each tool to perform the analysis.

For clustering, we compared scDHA with SC3, SEURAT, SINCERA, CIDR, and k-means. We used the following packages: i) SC3 version 1.10.1 from Bioconductor, ii) SEURAT version 2.3.4 from CRAN, iii) CIDR version 0.1.5 from github (github.com/VCCRI/CIDR), iv) SINCERA script provided by Hemberg group (scrnaseq-course.cog.sanger.ac.uk/website/biological-analysis.html), and v) stats for k-means in conjunction with PCA implementation available in the package irlba version 2.3.3 from CRAN. For k-means, we used the first 100 principal components for clustering purpose. In contrast to the other five methods, k-means cannot determine the number of clusters. Therefore, we also provided the true number of cell types for k-means. In addition, since k-means often converges to local optima, we ran k-means using 1,000 different sets of starting points and then chose the partitioning with the smallest squared error.

For dimension reduction and visualization, we used the following packages: i) irlba version 2.3.3 from CRAN for PCA, ii) Rtsne version 0.15 from CRAN for t-SNE, and iii) python package umap-learn version 0.3.9 from Anaconda python distribution for UMAP. UMAP python package is run through a wrapper in R package umap version 0.2.2.

For classification, we compared scDHA with XGBoost, Random Forest (RF), Deep Learning (DL), and Gradient Boosting Machine (GBM). We used the R package H2O version 3.24.0.5 from CRAN. This package provides the implementation of XGBoost, RF, DL, and GBM. All models were run with 5-fold cross validation for better accuracy.

For time-trajectory inference, we compared scDHA with Monocle, TSCAN, and Slingshot. We used the following packages: i) R package Monocle3 version 0.1.1 from github (github.com/cole-trapnell-lab/monocle3), ii) TSCAN version 1.20.0 from Bioconductor, and iii) Slingshot version 1.3.1 from Bioconductor.

### scDHA Pipeline

scDHA requires an expression matrix *M* as input, in which rows represent cells and columns represents genes/transcripts. scDHA pipeline for sc-RNA sequencing data analysis consists of two core modules (Figure 1a). The first module is a non-negative kernel autoencoder that provides a non-negative, part-based representation of the data. Based on the weight distribution of the encoder, scDHA removes genes or components that have insignificant contribution to the representation. The second module is a Stacked Bayesian Self-learning Network that is built upon the Variational Autoencoder^9^ to project the data onto a low dimensional space. In short, for example clustering application, the input data is normalized and insignificant genes are filtered to account for noises from technical variability. Processed data is then projected to low dimension latent space using a deep-learning approach and then clustered using k nearest neighbors spectral clustering. The detail of each step is described below.

### Data normalization and gene filtering

To reduce the technical variability coming from sequencing technologies for each cell, the expression data is normalized to range from 0 to 1 for each cell as follow:

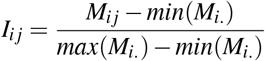

This normalization also helps speed up convergence of the projection model in the next step.

The normalized input data is then passed through an 1-layer autoencoder to filter out insignificant genes/features. Autoencoder is a self learning model, where the input data learn from itself. In short, autoencoder consists of two part: encoder and decoder. Data is processed through encoder to generate a much lower dimension data, and decoder uses this data to infer back the original data. Optimizing this process would enable output of the encoder to be used as a representation of original data. Based on the weights distribution of the encoder, genes with highest weight variance are selected for next step. High variability in weights means the gene contributes a meaningful information to some specific latent features, which is more useful for data representation. Moreover, using our strategy prevents the case of selecting genes with high variance but only within the same group of cells (highly correlated genes). Gene filtering removes the noise come from insignificant genes as well as speeds up the running time significantly.

### Stacked Bayesian Auto-encoder

A modified version of Variational Autoencoder (VAE, theorized by Kingma *et. al.*^9^), Stacked Bayesian Auto-encoder (Figure 4), is used as dimension reduction method. In brief, VAE model has the same basic structure as normal autoencoder, which is a self learning model, and consists of two components: encoder and decoder. Input data is processed using encoder to generate representative latent variables *z* with much smaller number of features compared to input. The original data then can be inferred from compressed data using decoder. By training the model to minimize the different between inferred and original data, the middle bottle neck layer can be viewed as the projections of input onto a low dimension space and used for other tasks such as clustering. In VAE case, instead of a deterministic *z* for each data point, VAE mapped input to distribution with means *μ* and variance *σ*^2^, *z* is sampled from this distribution (Figure 4), *z* ~ *N*(*μ*, *σ*^2^). By adding randomness in generating *z*, VAE can prevent overfitting case, where auto-encoder is big enough to map every single data point to *z* without learning generalized representation of data. The formulation of this architecture could be written like this:

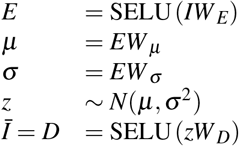

where E, D, represent encoder, decoder layer respectively. *μ*, *σ*, *z* are mean, variance and sampled latent layer. The input *I* is the filtered matrix from previous step. 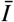 is the reconstruction of *I*.

**Figure 4.**
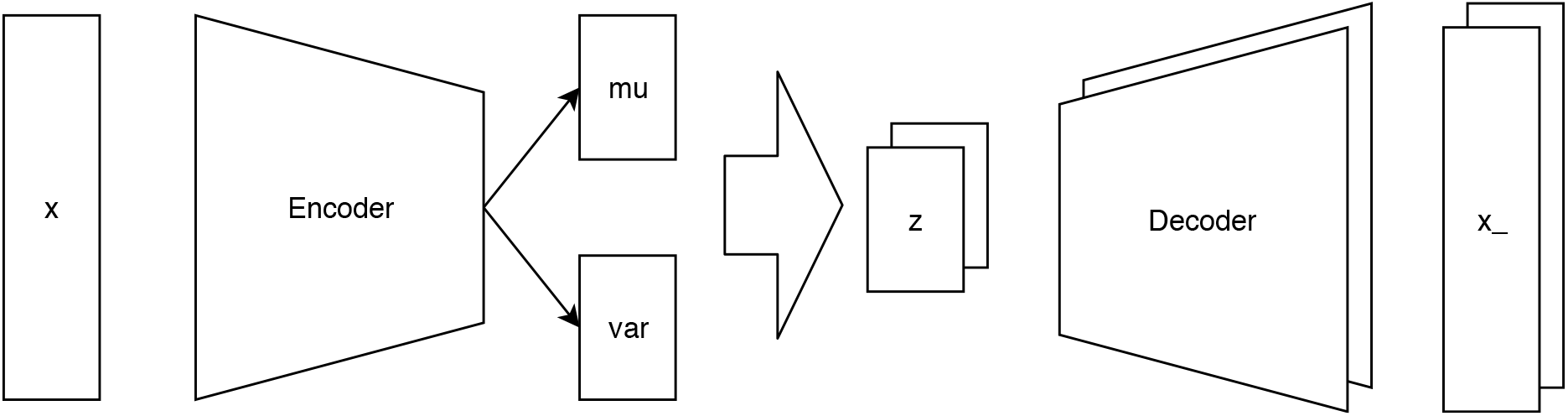
Stacked Bayesian Auto-encoder.

In our model, to further ensure the accuracy of inferred distribution, multiple latent spaces are sampled from Gaussian distributions with means *μ* and variances *σ*^2^ (Figure 4) using reparameterization trick,^9^ *z* = *μ* + *σ*∗*N*(0,1). The reparam-eterization trick is introduced to make sure that model can backpropagate, due to fact that *z* sampling process is non-differentiable and its gradient cannot be backpropagated. We keep the model size small to avoid over-fitting and force the network to learn the important features from the input data. Also, restricting the size of hidden layer will converge cells from the same group into similar latent space manifold. However, the size of the hidden layer needs to be sufficient to keep the latent variables disentangled.

To train our model, we used AdamW^53^ as optimizer and adapt two stage training scheme:^54^ (i) a *Warm-up* process which uses only reconstruction loss, and (ii) the VAE stage, in which the Kullback-Leibler loss is also considered to ensure the normal distribution of latent variables *z*. The warm-up process prevents model from ignoring reconstruction loss and only focusing on Kullback-Leibler loss which makes it fail to learn good representations of the individual data points. This process also helps the training of VAE model less sensitive to the initialized weights. For faster convergence speed and better accuracy, scaled exponential linear unit^55^ (SELU) is used as activation function. After finishing the training stage, the input data is processed through encoder to generate representative latent variables of original data.

### Cluster number prediction

The number of clusters is predicted based on two indices: (i) the ratio of “between sum of squares” over the “total sum of squares”:

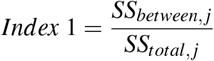

and (ii) the increase of total within the sum of squares when number of cluster increase:

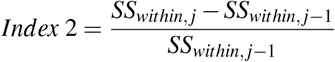

where *j* is number of cluster.

Bigger index 1 would means that members of one group are far from center of other groups, which means they are well separated. Index 2 is affected by the number of eigen-vectors generated by spectral decomposition, which is equal to number of cluster. We assumed that the addition of an eigenvector that leads to the highest spike in the total within sum of squares (which is undesirable) would be the number of clusters. These indices are calculated by performing k-nearest neighbor spectral clustering on a subset of samples over a range of cluster number. Mean of the predictions from these two indices is set to be the final number of cluster to input in clustering function.

### Clustering

k-nearest neighbor adaption of spectral clustering (k-nn SC) is selected as clustering method to improve the accuracy when dealing with non-spherical data distribution and still ensure the fast running time. However, instead of using Euclidean distance to determine the similarity between two samples, Pearson correlation is used to improve stability of cluster assignment. The different between k-nn SC and normal SC is that the constructed affinity matrix of data points is a sparse one with distance values for only k nearest neighbor of each point, the rest are zero. The clustering process of k-nn SC is consists of 4 steps: (i) constructing affinity matrix *A* for all data points to use as input graph, (ii) generating Symmetric normalized Laplacian matrix *L*^*sym*^ :=*I* − *D*^−½^*AD*^−½^ where *D* is the degree matrix of the graph, *A* is the constructed affinity matrix and *I* is identity matrix, (iii) calculating eigenvalues for Laplacian matrix and select ones with smallest values, generating eigenvectors corresponding to selected eigenvalues, performing final clustering using k-means on generated eigenvectors.

Because samples from the same group have been force to have the similar manifolds, it is not necessary to perform clustering on the entire dataset. Instead, for big dataset with more than 5,000 samples, we sampled randomly 2,000 cells and perform clustering on these, the rest of data was matched back to clustering assignment using votes from their nearest neighbors. This approach still ensures the high clustering quality without compromising the speed of method (Figure 5).

**Figure 5.**
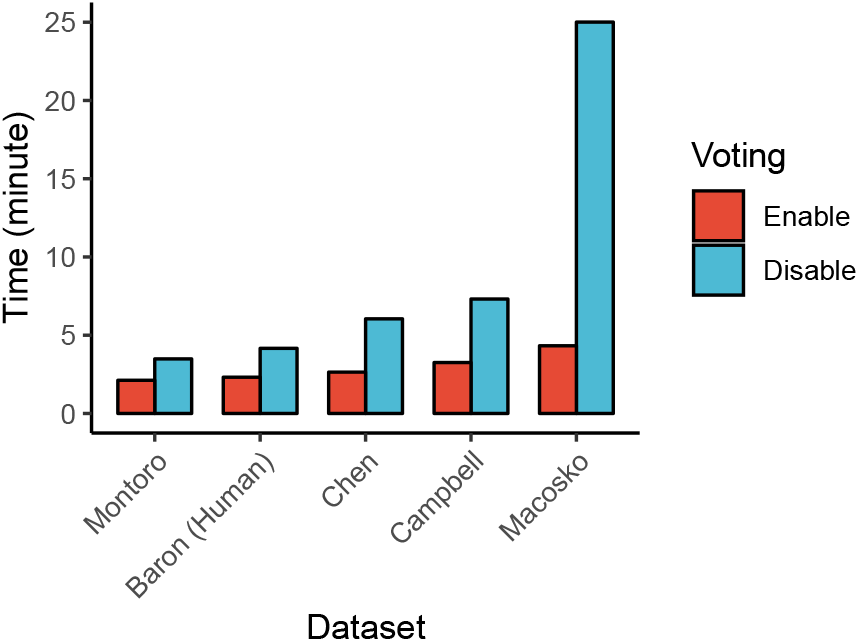
Running time of scDHA on big dataset with and without applying voting strategy.

### Consensus clustering

To achieve higher accuracy and preventing the situation when model converges to a local minimum, an ensemble of data projection models is used. Data projection and clustering process are repeated multiple times and each individual replicate generated different projected data. Then, cluster predictions from all replicate are combined using the Weighted-based meta-clustering (wMetaC) from package SHARP.^56^ wMetaC is conducted through 5 steps: (i) calculating cell-cell weighted similarity matrix *W*, *w*_*i,j*_ = *s*_*i,j*_(1 − *s*_*i,j*_) where *s*_*i,j*_ is the chance that cell *i* and *j* are in the same cluster, (ii) calculating cell weight, which is the sum of all cell-cell weights related to this cell, (iii) generating cluster-cluster similarity matrix |*C*|*x*|*C*|, where C is the union of all the clusters obtain in each individual replicates, (iv) performing hierarchical clustering on cluster-cluster similarity matrix, and (v) determining final results by voting scheme.

### Visualization

To get a better visualization, log and *z* transformations are applied to make the distribution of distances from one point to its neighbors more uniform:

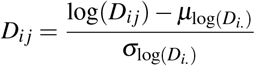

where *D* is a distance or similarity matrix between all samples, *D_i._* is row i-th.

After that, the probabilities *p_i j_* that proportionally to the similarity between sample *i* and *j* could be computed as follows:

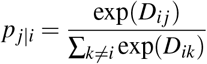

Our goal is to learn a 2-dimensional projection of data that retain probabilities *p* as well as possible. The same transformation as above would be applied to this projected data to generate probabilities *q*. We then minimize Kullback-Leibler divergence of distribution Q from P:

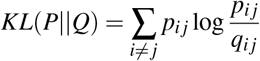

### Classification

We know that processing data using scDHA would make the biological relevant samples become more similar. Therefore, we can use the processed data to classify new data using available labeled data with better accuracy than using original data. By combining new and old data together using our processing pipeline, we can learn the underlying manifolds that can represent both datasets to improve classification accuracy and prevent overfitting when training the classifier. Classification using scDHA could be conducted through steps: (i) filtering new and labeled data using common genes from two datasets and then combine to 1 matrix, (ii) processing data using scDHA to get low dimension data, (iii) calculating distance from samples in new data to labeled data using Pearson distance, and (iv) assigning cluster for new datasets using group from labeled one using nearest neighbor classification.

### Time trajectory inference

To infer a pseudo time trajectory for single cell data, we use the compress data, output of our processing pipeline. Pearson distance between all samples are calculated to determine their similarities. We apply minimum spanning tree (MST) algorithm on the graph with sample as vertices and distances as edges to find the shortest path that goes through all the vertices. Because the denoising and dimension reduction process have already removed noise from the original data and made cells from the same group get closer to each others, applying MST algorithm on the graph would result to a path that pass through all data points and connect each cluster to its nearest neighbors. From this MST, pseudo time is determined by distance from one point to the designated starting point.

## Additional information

## Acknowledgments

This work was partially supported by the National Aeronautics and Space Administration (NASA) under Grant Number 80NSSC19M0170. Any opinions, findings, and conclusions or recommendations expressed in this material are those of the authors and do not necessarily reflect the views of any of the funding agencies.

## Author contributions

DT and TN conceived of and designed the approach. DT implemented the method in R, performed the data analysis and all computational experiments. BT and HN helped with data preparation and some data analysis. HL and CLV provided advice in method development. DT, HL and TN wrote the manuscript. All authors reviewed the manuscript.

