## Supplementary Material for "Fast and precise single-cell data analysis using hierarchical autoencoder"

Duc Tran[1], Hung Nguyen[1], Bang Tran[1], and Tin Nguyen[1,*]

[1] Department of Computer Science and Engineering, University of Nevada, Reno

### 1 Supplementary Note 1: Unsupervised clustering of single-cell data

#### 1.1 Evaluation metrics

We use three different metrics to assess the performance of the six clustering methods: Adjusted Rand Index (ARI), Normalized Mutual Information (NMI), and Jaccard index.

Rand index (RI)[1] measures the agreement between a given clustering and the ground truth. RI is calculated as:

$$RI = \frac{a+b}{a+b+c+d} = \frac{a+b}{\binom{N}{2}} \tag{1}$$

where $a$ is the number of pairs that belong to the same true cell type and are clustered together, $b$ is the number of pairs that belong to different true cell types and are not clustered together, $c$ is the number of pairs that belong to the same cell types and are not clustered together, $d$ is the number of pairs that belong to different cell types and are clustered together, and $\binom{N}{2}$ is the number of possible pairs that can be formed from the $N$ cells. Intuitively, RI is the fraction of pairs that are grouped in the same way (either together or not) in the two partitions compared (e.g. 0.9 means 90% of pairs are grouped in the same way). The Adjusted Rand Index (ARI)[2] is the corrected-for-chance version of the Rand Index. The ARI takes values from -1 to 1, with the ARI expected to be 0 for a random subtyping.

Jaccard index (JI) is also known as Intersection over Union. In our context, The Jaccard index is basically the number of pairs that belong to the same true cell type and are clustered together, divided by the number of pairs that are either in the same true cell type or are clustered together. JI is calculated as:

$$J = \frac{a}{a+b+c} \tag{2}$$

Finally, Normalized Mutual Information (NMI) is a normalized version of Mutual Information (MI). Denoting $X$ as the true labeling of the cells and $Y$ is the partitioning obtained from a clustering method, the NMI is calculated as:

$$NMI = \frac{1}{2} \times \frac{I(X;Y)}{H(X)+H(Y)} \tag{3}$$

where $I(X;Y)$ is the mutual information between $X$ and $Y$. $H(X)$ is the entropy of the true partition $X$ and $H(Y)$ is the entropy of the partition obtained from clustering. The NMI value take a range from 0 to 1 in which 1 indicates a perfect match between cell types and clusters. In contrast, 0 value means no mutual information between cell types and clusters.

#### 1.2 Performance evaluation

We assess the performance of the six clustering methods, scDHA, SC3,[3] SEURAT,[4] SINCERA,[5] CIDR,[6] and k-means, using ARI, NMI and JI described above. Table S1 shows the ARI values while Tables S2 and S3 shows the NMI and JI values, respectively. Regardless of the evaluation metrics, scDHA consistently outperforms other five methods.

**Table 1.** Performance of scDHA, SC3, SEURAT, SINCERA, CIDR, and k-means on 24 single cell datasets measured by ARI. Cells highlighted in green have the highest ARI values. The average ARI of scDHA is 0.83, which is much higher than the rest (0.5 is the second best). In addition, scDHA has the highest ARI values in all but one dataset.

| Dataset | #Sample | #Type | scDHA | SC3 | SEURAT | SINCERA | CIDR | k-means |
|---|---|---|---|---|---|---|---|---|
| 1. Yan | 90 | 6 | 0.86 | 0.66 | 0.39 | 0.72 | 0.80 | 0.80 |
| 2. Goolam | 124 | 5 | 0.84 | 0.60 | 0.42 | 0.30 | 0.70 | 0.48 |
| 3. Deng | 268 | 6 | 0.89 | 0.44 | 0.29 | 0.70 | 0.51 | 0.60 |
| 4. Pollen | 301 | 11 | 0.92 | 0.96 | 0.61 | 0.85 | 0.90 | 0.89 |
| 5. Patel | 430 | 5 | 0.87 | 0.46 | 0.76 | 0.47 | 0.45 | 0.82 |
| 6. Wang | 457 | 7 | 0.85 | 0.85 | 0.65 | 0.29 | 0.63 | 0.44 |
| 7. Darmanis | 466 | 9 | 0.68 | 0.44 | 0.58 | 0.55 | 0.50 | 0.44 |
| 8. Camp (Brain) | 553 | 5 | 0.86 | 0.56 | 0.65 | 0.59 | 0.34 | 0.48 |
| 9. Usoskin | 622 | 4 | 0.82 | 0.80 | 0.66 | 0.38 | 0.82 | 0.23 |
| 10. Kolodziejczyk | 704 | 3 | 0.87 | 0.44 | 0.45 | 0.46 | 0.43 | 0.48 |
| 11. Camp (Liver) | 777 | 7 | 0.74 | 0.66 | 0.71 | 0.49 | 0.61 | 0.53 |
| 12. Xin | 1,600 | 8 | 0.95 | 0.14 | 0.42 | 0.16 | 0.57 | 0.44 |
| 13. Baron (Mouse) | 1,886 | 13 | 0.88 | 0.26 | 0.49 | 0.39 | 0.47 | 0.29 |
| 14. Muraro | 2,126 | 10 | 0.91 | 0.38 | 0.57 | 0.32 | 0.22 | 0.34 |
| 15. Segerstolpe | 2,209 | 14 | 0.92 | 0.29 | 0.44 | 0.40 | 0.37 | 0.29 |
| 16. Klein | 2,717 | 4 | 0.98 | 0.45 | 0.54 | 0.61 | 0.68 | 0.29 |
| 17. Romanov | 2,881 | 7 | 0.75 | 0.22 | 0.39 | 0.23 | 0.32 | 0.30 |
| 18. Zeisel | 3,005 | 9 | 0.80 | 0.33 | 0.51 | 0.42 | 0.37 | 0.36 |
| 19. Lake | 3,042 | 16 | 0.60 | 0.39 | 0.48 | 0.31 | 0.47 | 0.38 |
| 20. Montoro | 7,193 | 7 | 0.81 | 0.11 | 0.24 | 0.13 | 0.30 | 0.45 |
| 21. Baron (Human) | 8,569 | 14 | 0.93 | 0.14 | 0.58 | 0.34 | 0.73 | 0.41 |
| 22. Chen | 12,089 | 46 | 0.78 | 0.16 | 0.63 | 0.60 | 0.36 | 0.33 |
| 23. Campbell | 21,086 | 21 | 0.64 | 0.07 | 0.37 | 0.00 | 0.23 | 0.13 |
| 24. Macosko | 44,808 | 12 | 0.73 | 0.07 | 0.22 | 0.41 | 0.17 | 0.25 |
| Mean ARI | | | 0.83 | 0.41 | 0.50 | 0.42 | 0.50 | 0.44 |

**Table 2.** Performance of scDHA, SC3, SEURAT, SINCERA, CIDR, and k-means on 24 single cell datasets measured by NMI. Cells highlighted in green have the highest NMI values. scDHA outperforms other methods by having the highest average NMI value. In addition, scDHA has the highest NMI values in 21 out of 24 datasets.

| Dataset | #Sample | #Type | scDHA | SC3 | SEURAT | SINCERA | CIDR | k-means |
|---|---|---|---|---|---|---|---|---|
| 1. Yan | 90 | 6 | 0.89 | 0.80 | 0.55 | 0.82 | 0.84 | 0.86 |
| 2. Goolam | 124 | 5 | 0.82 | 0.80 | 0.61 | 0.61 | 0.78 | 0.63 |
| 3. Deng | 268 | 6 | 0.89 | 0.73 | 0.53 | 0.73 | 0.74 | 0.78 |
| 4. Pollen | 301 | 11 | 0.96 | 0.95 | 0.80 | 0.93 | 0.94 | 0.94 |
| 5. Patel | 430 | 5 | 0.84 | 0.67 | 0.76 | 0.67 | 0.57 | 0.83 |
| 6. Wang | 457 | 7 | 0.83 | 0.81 | 0.71 | 0.43 | 0.71 | 0.57 |
| 7. Darmanis | 466 | 9 | 0.75 | 0.67 | 0.64 | 0.66 | 0.64 | 0.62 |
| 8. Camp (Brain) | 553 | 5 | 0.82 | 0.68 | 0.70 | 0.62 | 0.49 | 0.55 |
| 9. Usoskin | 622 | 4 | 0.81 | 0.79 | 0.74 | 0.54 | 0.80 | 0.31 |
| 10. Kolodziejczyk | 704 | 3 | 0.90 | 0.68 | 0.68 | 0.54 | 0.57 | 0.51 |
| 11. Camp (Liver) | 777 | 7 | 0.85 | 0.81 | 0.85 | 0.69 | 0.79 | 0.72 |
| 12. Xin | 1,600 | 8 | 0.87 | 0.39 | 0.60 | 0.42 | 0.55 | 0.60 |
| 13. Baron (Mouse) | 1,886 | 13 | 0.85 | 0.65 | 0.75 | 0.61 | 0.51 | 0.59 |
| 14. Muraro | 2,126 | 10 | 0.88 | 0.69 | 0.77 | 0.51 | 0.43 | 0.53 |
| 15. Segerstolpe | 2,209 | 14 | 0.90 | 0.65 | 0.75 | 0.62 | 0.45 | 0.53 |
| 16. Klein | 2,717 | 4 | 0.97 | 0.69 | 0.71 | 0.67 | 0.66 | 0.40 |
| 17. Romanov | 2,881 | 7 | 0.69 | 0.43 | 0.60 | 0.31 | 0.34 | 0.35 |
| 18. Zeisel | 3,005 | 9 | 0.78 | 0.62 | 0.67 | 0.47 | 0.47 | 0.55 |
| 19. Lake | 3,042 | 16 | 0.67 | 0.68 | 0.73 | 0.47 | 0.54 | 0.62 |
| 20. Montoro | 7,193 | 7 | 0.74 | 0.30 | 0.50 | 0.24 | 0.46 | 0.56 |
| 21. Baron (Human) | 8,569 | 14 | 0.88 | 0.50 | 0.80 | 0.46 | 0.72 | 0.63 |
| 22. Chen | 12,089 | 46 | 0.77 | 0.53 | 0.79 | 0.53 | 0.42 | 0.63 |
| 23. Campbell | 21,086 | 21 | 0.68 | 0.49 | 0.74 | 0.15 | 0.38 | 0.49 |
| 24. Macosko | 44,808 | 12 | 0.59 | 0.31 | 0.56 | 0.19 | 0.33 | 0.40 |
| Mean NMI | | | 0.82 | 0.64 | 0.69 | 0.54 | 0.59 | 0.59 |

**Table 3.** Performance of scDHA, SC3, SEURAT, SINCERA, CIDR, and k-means on 24 single cell datasets measured by Jaccard Index (JI). Cells highlighted in green have the highest JI values. scDHA outperforms other methods by having the highest average JI value. scDHA also has the highest JI values in 22 out of 24 datasets.

| Dataset | #Sample | #Type | scDHA | SC3 | SEURAT | SINCERA | CIDR | k-means |
|---|---|---|---|---|---|---|---|---|
| 1. Yan | 90 | 6 | 0.80 | 0.57 | 0.38 | 0.64 | 0.73 | 0.73 |
| 2. Goolam | 124 | 5 | 0.82 | 0.54 | 0.46 | 0.28 | 0.65 | 0.45 |
| 3. Deng | 268 | 6 | 0.86 | 0.40 | 0.33 | 0.65 | 0.46 | 0.55 |
| 4. Pollen | 301 | 11 | 0.87 | 0.93 | 0.50 | 0.76 | 0.83 | 0.82 |
| 5. Patel | 430 | 5 | 0.81 | 0.35 | 0.67 | 0.36 | 0.38 | 0.75 |
| 6. Wang | 457 | 7 | 0.81 | 0.81 | 0.58 | 0.31 | 0.56 | 0.39 |
| 7. Darmanis | 466 | 9 | 0.59 | 0.34 | 0.48 | 0.46 | 0.42 | 0.35 |
| 8. Camp (Brain) | 553 | 5 | 0.82 | 0.48 | 0.57 | 0.55 | 0.30 | 0.44 |
| 9. Usoskin | 622 | 4 | 0.76 | 0.74 | 0.58 | 0.35 | 0.78 | 0.29 |
| 10. Kolodziejczyk | 704 | 3 | 0.83 | 0.37 | 0.38 | 0.44 | 0.40 | 0.50 |
| 11. Camp (Liver) | 777 | 7 | 0.64 | 0.54 | 0.59 | 0.40 | 0.49 | 0.44 |
| 12. Xin | 1,600 | 8 | 0.94 | 0.13 | 0.39 | 0.15 | 0.58 | 0.41 |
| 13. Baron (Mouse) | 1,886 | 13 | 0.85 | 0.20 | 0.41 | 0.34 | 0.42 | 0.25 |
| 14. Muraro | 2,126 | 10 | 0.87 | 0.29 | 0.46 | 0.32 | 0.23 | 0.30 |
| 15. Segerstolpe | 2,209 | 14 | 0.88 | 0.21 | 0.34 | 0.35 | 0.35 | 0.24 |
| 16. Klein | 2,717 | 4 | 0.97 | 0.36 | 0.46 | 0.58 | 0.61 | 0.33 |
| 17. Romanov | 2,881 | 7 | 0.69 | 0.17 | 0.31 | 0.29 | 0.31 | 0.28 |
| 18. Zeisel | 3,005 | 9 | 0.73 | 0.24 | 0.41 | 0.41 | 0.37 | 0.30 |
| 19. Lake | 3,042 | 16 | 0.52 | 0.28 | 0.37 | 0.27 | 0.39 | 0.29 |
| 20. Montoro | 7,193 | 7 | 0.80 | 0.11 | 0.23 | 0.13 | 0.29 | 0.43 |
| 21. Baron (Human) | 8,569 | 14 | 0.89 | 0.10 | 0.46 | 0.29 | 0.65 | 0.32 |
| 22. Chen | 12,089 | 46 | 0.68 | 0.11 | 0.51 | 0.49 | 0.29 | 0.23 |
| 23. Campbell | 21,086 | 21 | 0.62 | 0.06 | 0.30 | 0.17 | 0.35 | 0.11 |
| 24. Macosko | 44,808 | 12 | 0.76 | 0.08 | 0.22 | 0.50 | 0.24 | 0.28 |
| Mean Jaccard Index | | | 0.78 | 0.35 | 0.43 | 0.40 | 0.46 | 0.40 |

**Table 4.** Running time of scDHA, SC3, SEURAT, SINCERA, CIDR, and k-means on 24 single cell datasets. Overall, scDHA is the fastest with an average running time of 2 minute per dataset.

| Dataset | scDHA | SC3 | SEURAT | SINCERA | CIDR | k-means |
|---|---|---|---|---|---|---|
| Yan | 1.12 | 0.5 | 0.89 | 0.03 | 0.02 | 0.03 |
| Goolam | 1.43 | 0.53 | 1.02 | 0.06 | 0.04 | 0.12 |
| Deng | 1.45 | 0.6 | 1.07 | 0.07 | 0.07 | 0.39 |
| Pollen | 1.61 | 0.66 | 1.22 | 0.08 | 0.08 | 0.5 |
| Patel | 1.2 | 0.98 | 0.86 | 0.04 | 0.04 | 0.19 |
| Wang | 1.51 | 0.95 | 1.5 | 0.1 | 0.12 | 0.76 |
| Darmanis | 1.63 | 0.99 | 1.29 | 0.11 | 0.12 | 0.7 |
| Camp (Brain) | 1.62 | 1.18 | 1.32 | 0.12 | 0.12 | 0.75 |
| Usoskin | 1.66 | 1.36 | 1.19 | 0.18 | 0.18 | 1.13 |
| Kolodziejczyk | 1.92 | 1.57 | 1.81 | 0.31 | 0.32 | 1.82 |
| Camp (Liver) | 2.07 | 1.82 | 1.2 | 0.19 | 0.25 | 0.83 |
| Xin | 2.43 | 15.11 | 2.87 | 1.3 | 1.2 | 3.35 |
| Baron (Mouse) | 2.43 | 18.25 | 1.42 | 0.68 | 0.78 | 1.25 |
| Muraro | 2.68 | 3.99 | 1.42 | 1.09 | 1.31 | 1.51 |
| Segerstolpe | 2.71 | 4.34 | 1.9 | 1.56 | 1.89 | 2.58 |
| Klein | 2.56 | 9.46 | 3.66 | 2.2 | 2.37 | 4.11 |
| Romanov | 2.57 | 8.34 | 2.07 | 2.45 | 2.65 | 2.97 |
| Zeisel | 2.57 | 9.13 | 2.25 | 2.18 | 2.72 | 1.92 |
| Lake | 2.61 | 10.59 | 3.73 | 2.8 | 3.88 | 3.91 |
| Montoro | 2.62 | 60.4 | 2.82 | 17.19 | 15.95 | 8.86 |
| Baron (Human) | 2.82 | 52.22 | 3.05 | 17.45 | 33.3 | 5.71 |
| Chen | 3.14 | 68.93 | 4.76 | 40.32 | 84.39 | 13.26 |
| Campbell | 3.83 | 83.32 | 8.14 | 142.22 | 367 | 28.58 |
| Macosko | 4.7 | 163.45 | 27.36 | 571.45 | 3035.84 | 64.93 |
| Mean | 2.29 | 21.61 | 3.28 | 33.51 | 148.11 | 6.26 |

#### 2 Supplementary Note 2: Dimension reduction and visualization

We compare scDHA with PCA, t-SNE,[7,8] and UMAP.[9,10] Table S5 shows the silhouette index of the 24 dataset using each method while Table S6 shows the running time. The color-coded representations are shown in Figures S1–S6. In each representation, we calculate the silhouette index that measures the cohesion among the cells of the same type and the separation between different cell types. scDHA offers a significant improvement over current state-of-the-art techniques.

**Table 5.** Silhouette index calculated for each representation using scDHA, PCA, t-SNE, and UMAP. Cells highlighted in green have the highest silhouette values. scDHA has the highest average silhouette value. It also outperforms other methods in 21 out of 24 datasets.

| Dataset | scDHA | PCA | t-SNE | UMAP |
|---|---|---|---|---|
| Yan | 0.52 | 0.54 | 0.45 | 0.47 |
| Goolam | 0.37 | 0.31 | 0.28 | 0.27 |
| Deng | 0.5 | 0.6 | 0.49 | 0.67 |
| Pollen | 0.78 | 0.3 | 0.61 | 0.58 |
| Patel | 0.62 | 0.17 | 0.52 | 0.52 |
| Wang | 0.28 | -0.07 | 0.13 | 0.21 |
| Darmanis | 0.47 | 0.01 | 0.31 | 0.34 |
| Camp (Brain) | 0.54 | 0.07 | 0.36 | 0.3 |
| Usoskin | 0.62 | 0.07 | 0.4 | 0.51 |
| Kolodziejczyk | 0.81 | 0.3 | 0.43 | 0.54 |
| Camp (Liver) | 0.67 | 0.17 | 0.42 | 0.5 |
| Xin | 0.67 | 0.08 | 0.25 | 0.17 |
| Baron (Mouse) | 0.44 | -0.23 | 0.05 | 0.1 |
| Muraro | 0.57 | -0.2 | 0.24 | 0.46 |
| Segerstolpe | 0.66 | -0.22 | 0.01 | 0.24 |
| Klein | 0.72 | 0.24 | 0.48 | 0.69 |
| Romanov | 0.37 | 0.03 | 0.24 | 0.34 |
| Zeisel | 0.67 | 0.03 | 0.31 | 0.55 |
| Lake | 0.35 | -0.11 | 0.25 | 0.32 |
| Montoro | 0.16 | 0.24 | 0.09 | 0.29 |
| Baron (Human) | 0.61 | -0.14 | 0.2 | 0.46 |
| Chen | 0.49 | -0.07 | 0.09 | 0.35 |
| Campbell | 0.01 | -0.31 | -0.05 | -0.08 |
| Macosko | 0.27 | 0.11 | 0.09 | 0.36 |
| Mean | 0.51 | 0.08 | 0.28 | 0.38 |

**Table 6.** Running time of scDHA, PCA, t-SNE, and UMAP on 24 single cell datasets.

| Dataset | scDHA | PCA | t-SNE | UMAP |
|---|---|---|---|---|
| Yan | 1.43 | 0 | 0.02 | 0 |
| Goolam | 1.74 | 0.01 | 0.1 | 0.17 |
| Deng | 1.75 | 0.01 | 0.09 | 0.39 |
| Pollen | 1.87 | 0.02 | 0.13 | 0.41 |
| Patel | 1.43 | 0.02 | 0.07 | 0.17 |
| Wang | 1.75 | 0.02 | 0.2 | 0.67 |
| Darmanis | 1.9 | 0.02 | 0.21 | 0.67 |
| Camp (Brain) | 1.85 | 0.02 | 0.25 | 0.67 |
| Usoskin | 1.88 | 0.02 | 0.33 | 1.09 |
| Kolodziejczyk | 2.11 | 0.05 | 0.56 | 1.82 |
| Camp (Liver) | 2.08 | 0.02 | 0.14 | 0.67 |
| Xin | 2.7 | 0.1 | 0.98 | 2.9 |
| Baron (Mouse) | 2.72 | 0.04 | 0.29 | 0.98 |
| Muraro | 2.93 | 0.07 | 0.31 | 1.36 |
| Segerstolpe | 2.85 | 0.09 | 0.72 | 1.98 |
| Klein | 2.9 | 0.1 | 1.4 | 3.48 |
| Romanov | 2.99 | 0.1 | 0.53 | 2.68 |
| Zeisel | 2.91 | 0.08 | 0.37 | 1.43 |
| Lake | 3 | 0.11 | 0.67 | 3.38 |
| Montoro | 4.15 | 0.29 | 2.65 | 6.38 |
| Baron (Human) | 5.22 | 0.29 | 1.18 | 3.33 |
| Chen | 5.92 | 0.47 | 1.99 | 7.18 |
| Campbell | 6.64 | 1.07 | 4.18 | 10.93 |
| Macosko | 7.66 | 2.09 | 12.49 | 24.11 |
| Mean | 3.02 | 0.21 | 1.24 | 3.20 |

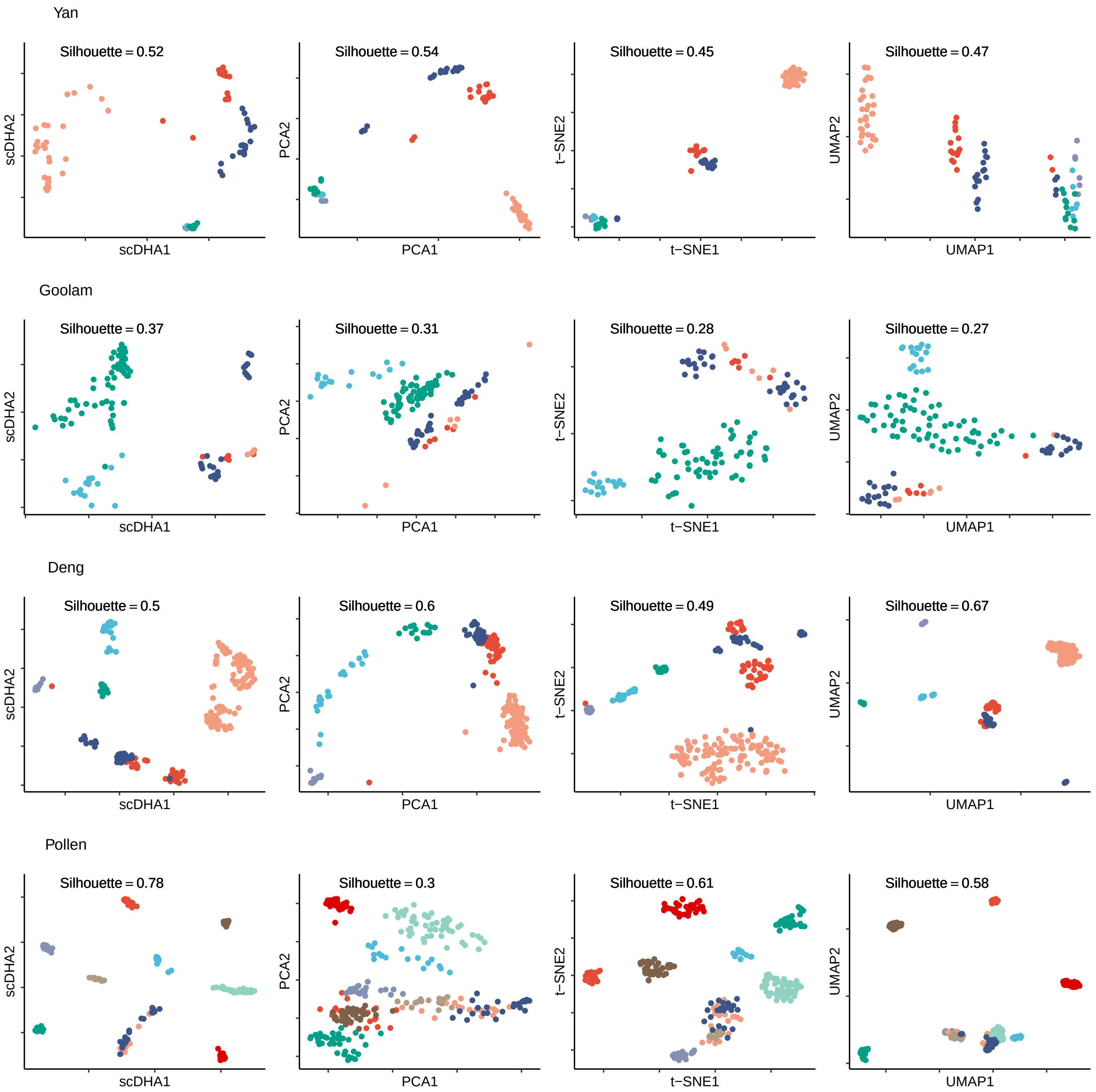


**Figure 1.** Representation of the Yan, Gollam, Deng, and Pollen datasets (top to bottom) using PCA, t-SNE, UMAP and scDHA (left to right). Different colors code for different cell types.

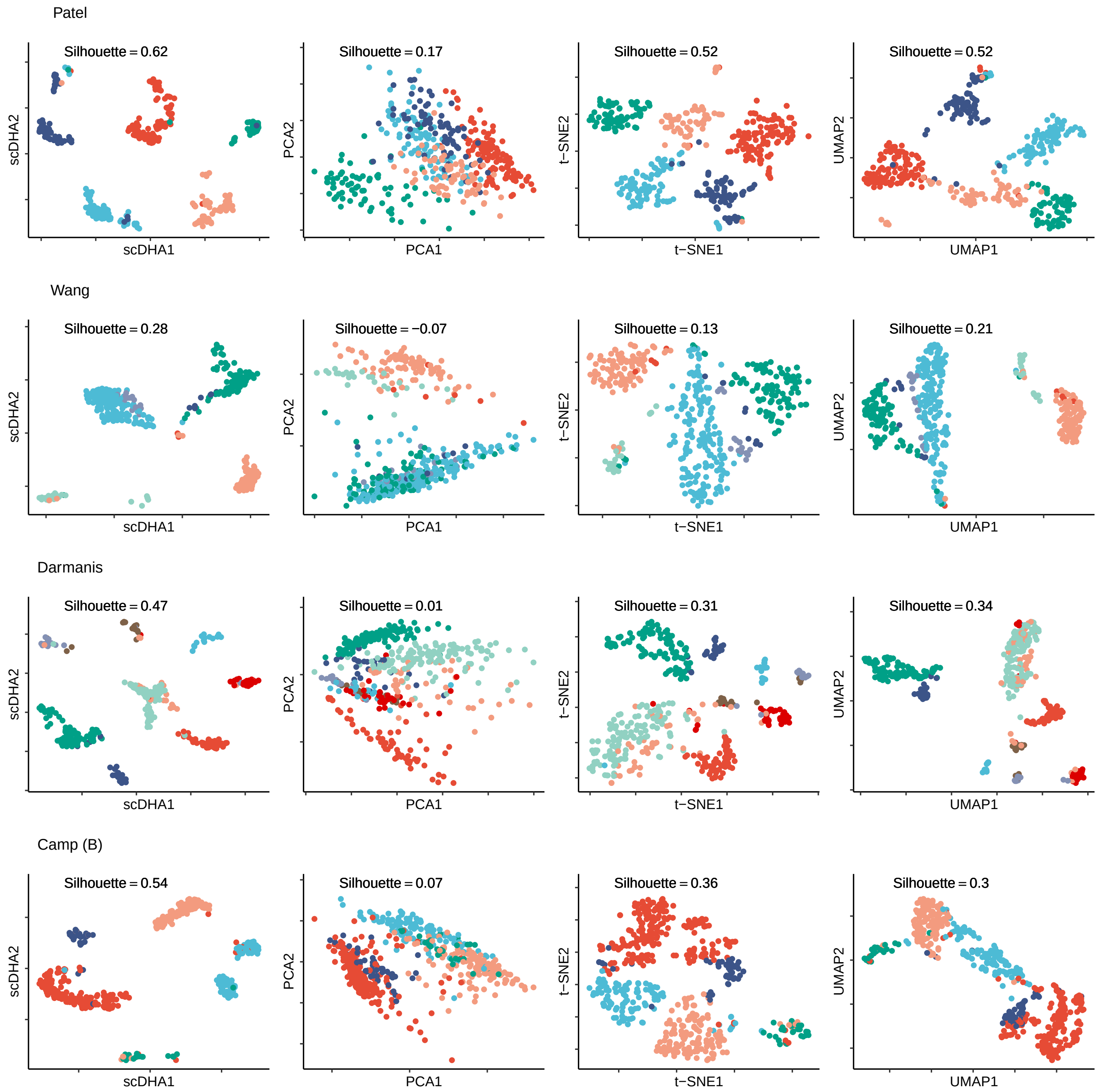


**Figure 2.** Representation of the Patel, Wang, Darmanis, and Camp (Brain) datasets (top to bottom) using PCA, t-SNE, UMAP and scDHA (left to right). Different colors code for different cell types.

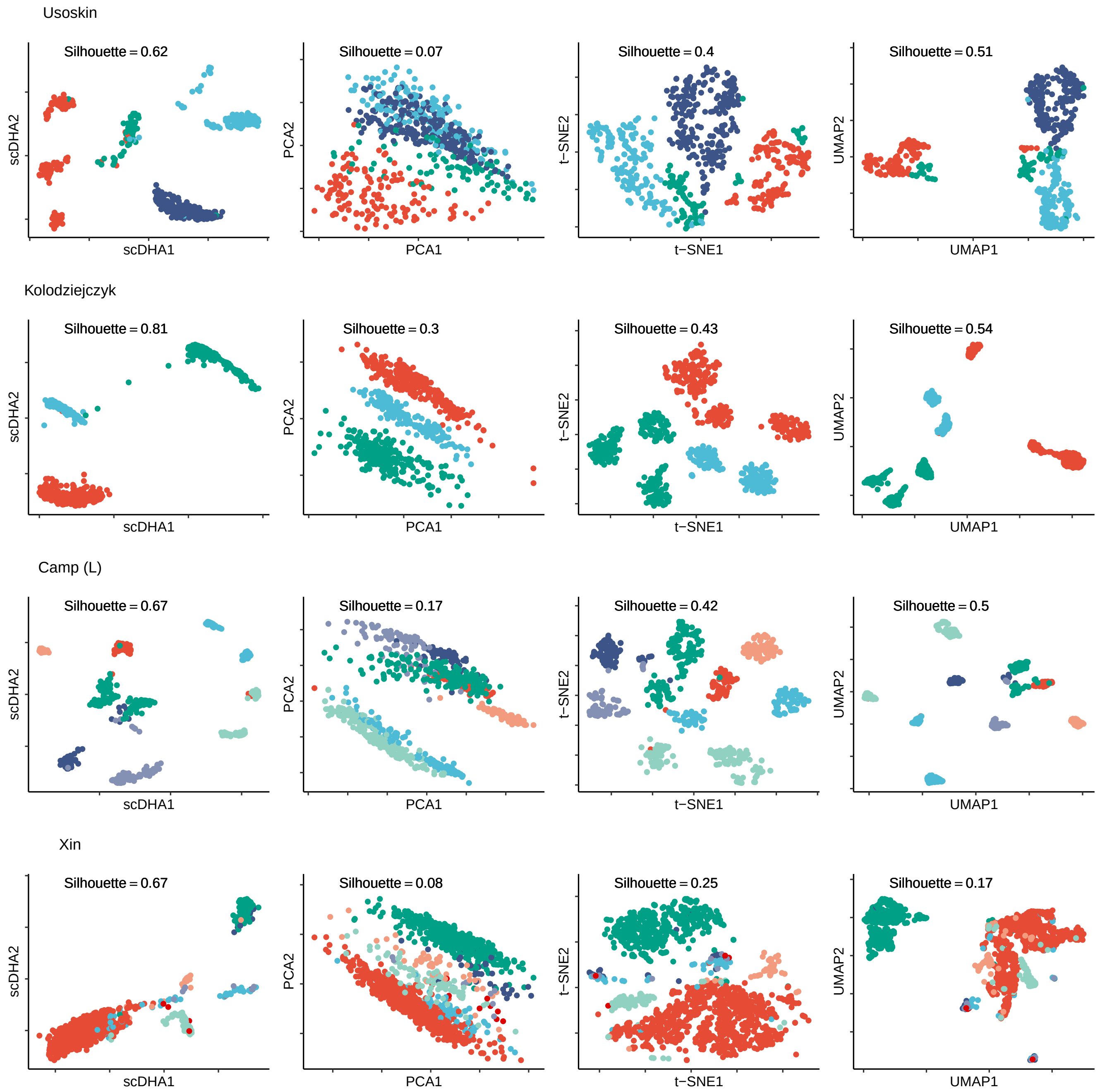


**Figure 3.** Representation of Usoskin, Kolodziejczyk, Camp (Liver), and Xin datasets (top to bottom) using PCA, t-SNE, UMAP and scDHA (left to right). Different colors code for different cell types.

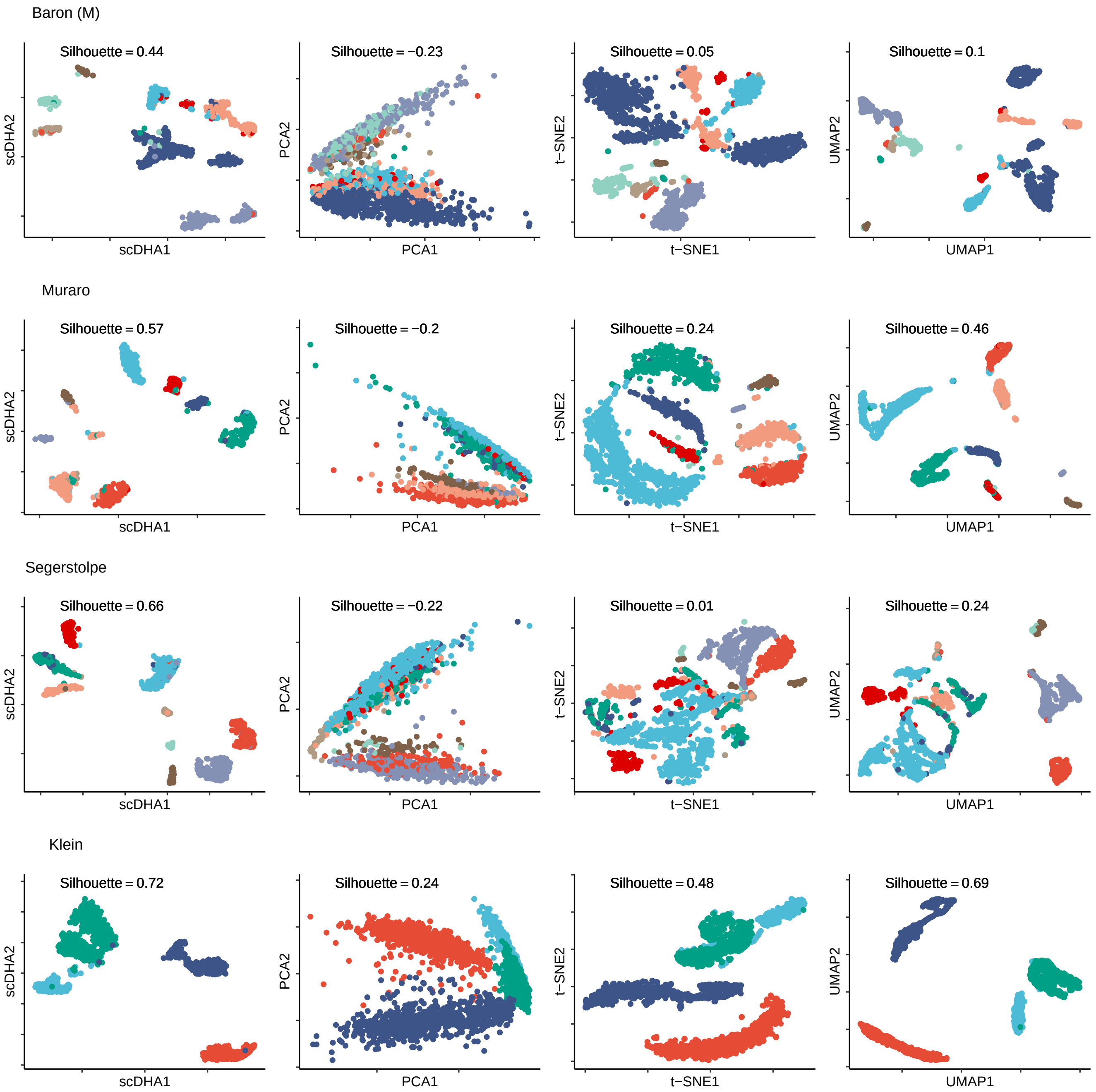


**Figure 4.** Representation of Baron (mouse), Muraro, Segerstolpe, and Klein datasets (top to bottom) using PCA, t-SNE, UMAP and scDHA (left to right). Different colors code for different cell types.

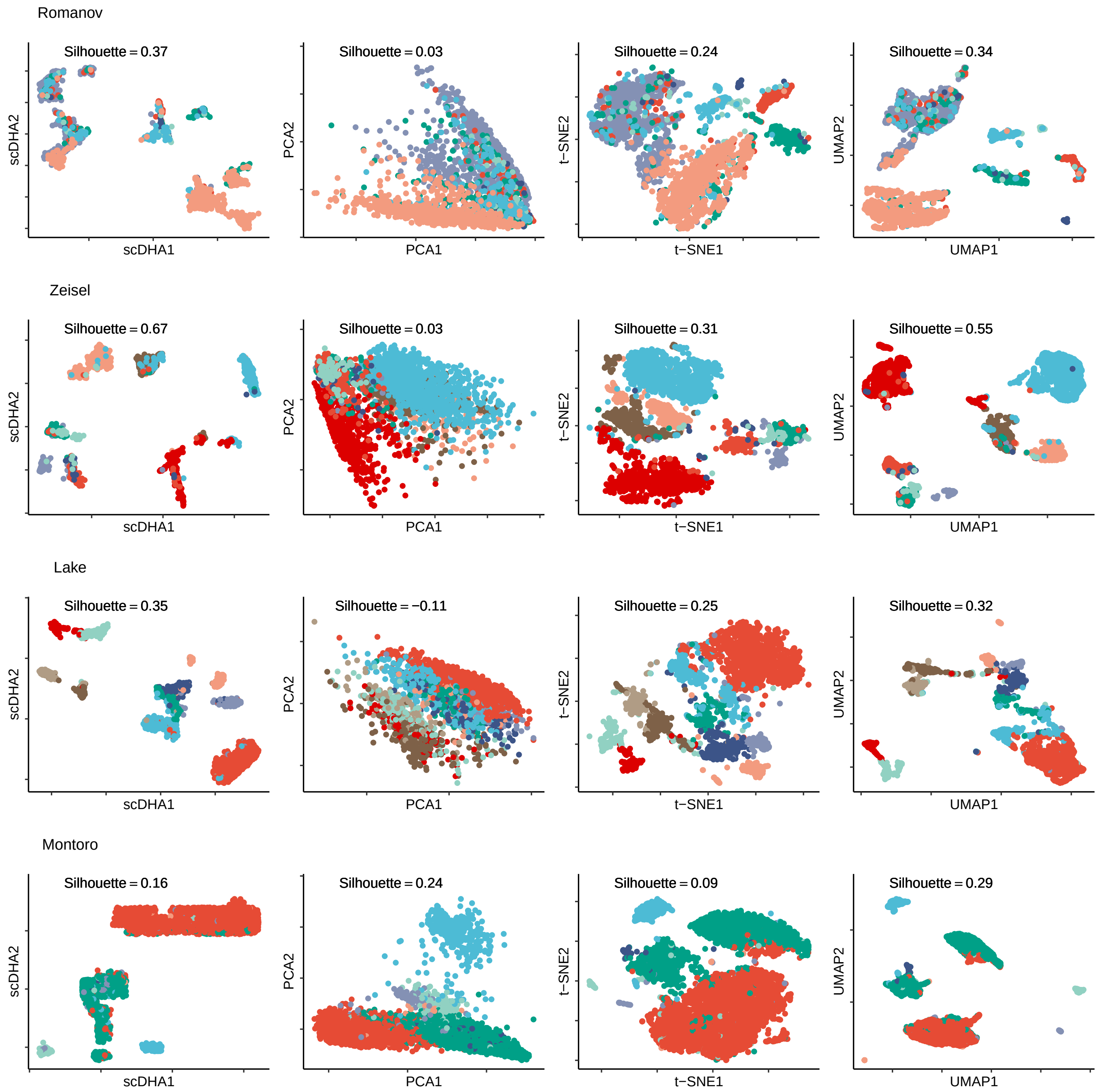


**Figure 5.** Representation of Romanov, Zeisel, Lake, and Montoro datasets (top to bottom) using PCA, t-SNE, UMAP and scDHA (left to right). Different colors code for different cell types.

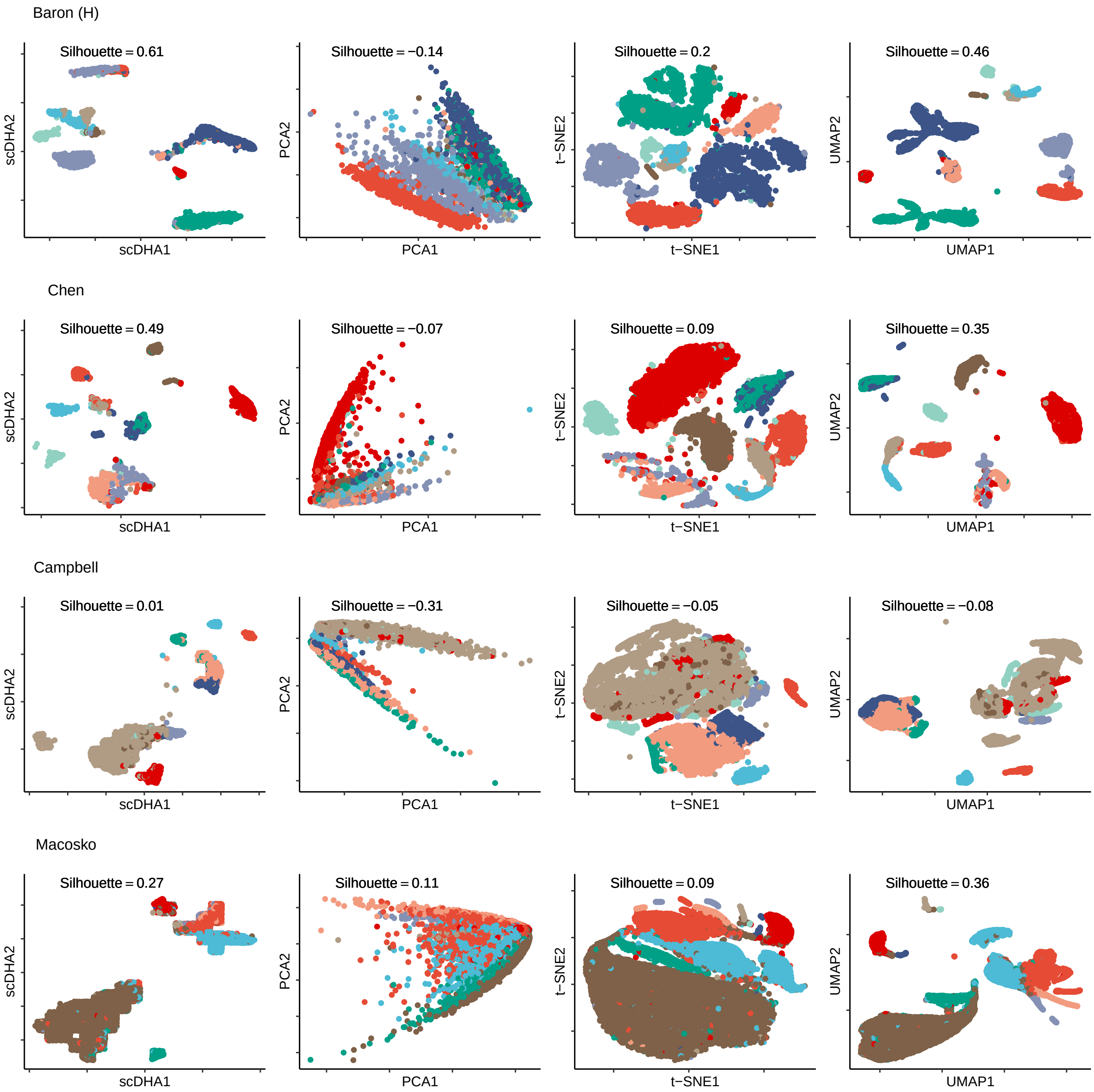


**Figure 6.** Representation of Baron (Human), Chen, Campbell, and Macosko datasets (top to bottom) using PCA, t-SNE, UMAP and scDHA (left to right). Different colors code for different cell types.

### 3 Supplementary Note 3: Cell classification

In this work, we compare scDHA with XGBoost,[11] Random Forest (RF),[12] Deep Learning (DL),[13] and Gradient Boosting Machine (GBM).[14] We use five datasets related to human pancreas to test the five classification methods. To calculate accuracy metric for classification comparison, we divide number of correct predictions to total number of samples. The accuracy for each evaluation sets is reported in the table below. Moreover, as seen in Figure S7, the running of our approach is much faster than other methods in comparison.

**Table 7.** Classification performance measuring by accuracy of scDHA, XGBoost, Random Forest (RF), Deep Learning (DL), and Gradient Boosting Machine (GBM) approach on single cell evaluation pairs.

| Training Dataset | Predicting Dataset | scDHA | XGBoost | RF | DL | GBM |
|---|---|---|---|---|---|---|
| Baron (Human) | Segerstolpe | 0.93 | 0.82 | 0.32 | 0.60 | 0.39 |
| Baron (Human) | Muraro | 0.88 | 0.86 | 0.79 | 0.72 | 0.74 |
| Baron (Human) | Xin | 0.99 | 0.93 | 0.49 | 0.03 | 0.84 |
| Baron (Human) | Wang | 0.96 | 0.27 | 0.28 | 0.01 | 0.60 |
| Segerstolpe | Baron (Human) | 0.94 | 0.83 | 0.71 | 0.21 | 0.49 |
| Segerstolpe | Muraro | 0.96 | 0.81 | 0.88 | 0.73 | 0.74 |
| Segerstolpe | Xin | 0.99 | 1.00 | 0.97 | 0.46 | 0.99 |
| Segerstolpe | Wang | 0.99 | 0.98 | 0.93 | 0.22 | 0.97 |
| Xin | Baron (Human) | 0.99 | 0.55 | 0.60 | 0.77 | 0.46 |
| Xin | Segerstolpe | 0.99 | 0.98 | 0.91 | 0.78 | 0.92 |
| Xin | Muraro | 0.97 | 0.70 | 0.82 | 0.57 | 0.42 |
| Xin | Wang | 1.00 | 1.00 | 0.58 | 0.58 | 0.96 |
| Muraro | Baron (Human) | 0.93 | 0.86 | 0.78 | 0.16 | 0.85 |
| Muraro | Segerstolpe | 0.97 | 0.93 | 0.65 | 0.65 | 0.72 |
| Muraro | Xin | 0.99 | 0.88 | 0.89 | 0.06 | 0.84 |
| Muraro | Wang | 0.98 | 0.85 | 0.64 | 0.01 | 0.73 |
| Wang | Baron (Human) | 0.93 | 0.14 | 0.38 | 0.30 | 0.38 |
| Wang | Segerstolpe | 0.92 | 0.90 | 0.75 | 0.44 | 0.91 |
| Wang | Muraro | 0.89 | 0.13 | 0.55 | 0.46 | 0.52 |
| Wang | Xin | 0.97 | 1.00 | 0.90 | 0.76 | 0.96 |

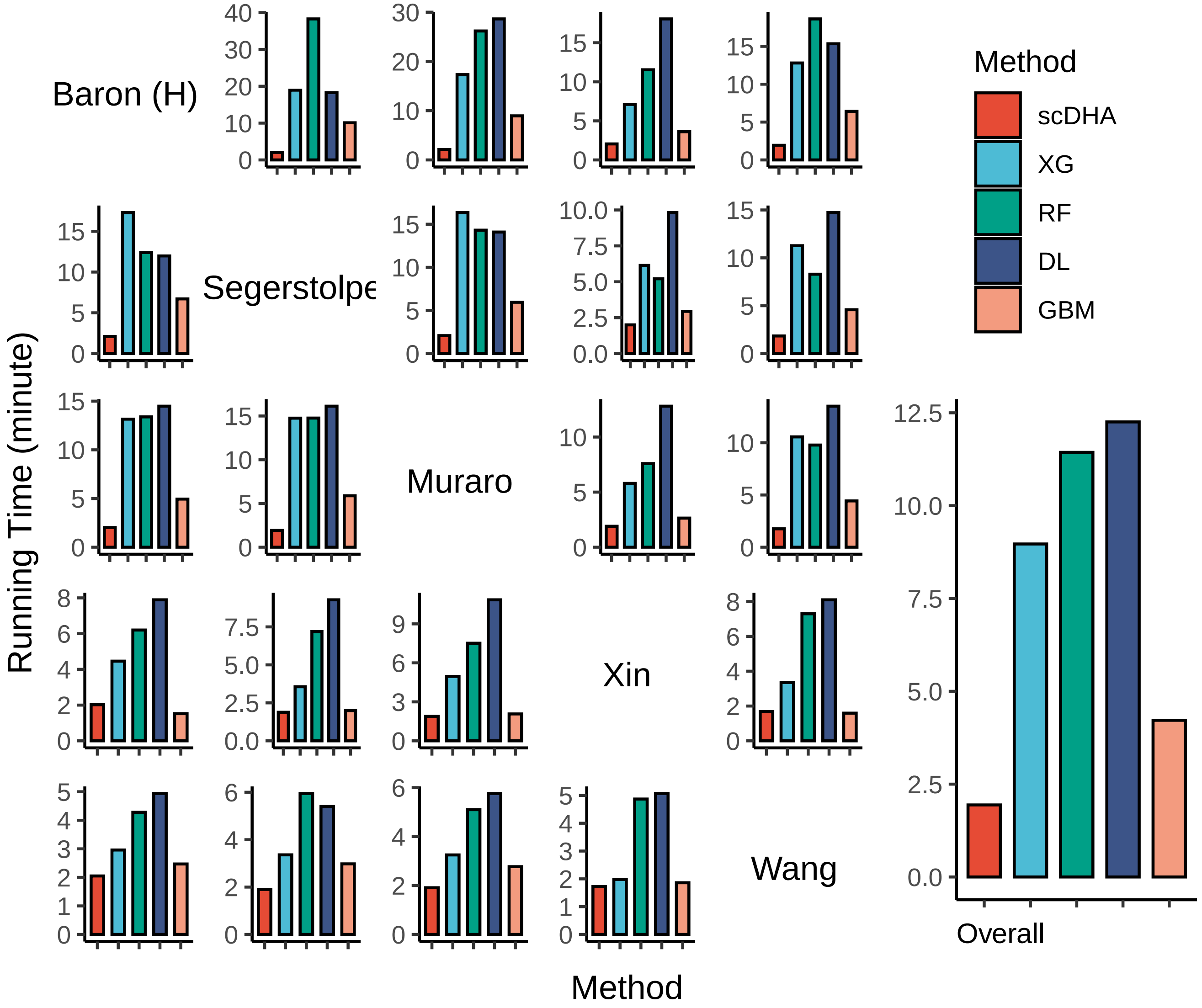


**Figure 7.** Running time of scDHA, XGBoost, Random Forest, Deep Learning and Gradient Boosting Machine approach on single cell evaluation pairs.

### 4 Supplementary Note 4: Time trajectory inference

We compare scDHA with Monocle,[15] TSCAN,[16] and Slingshot.[17] The pseudo-temporal ordering of the three mouse embryo datasets, Yan, Goolam, and Deng, are shown in Figure S8. The time trajectories inferred by each method are shown in Figure S9.

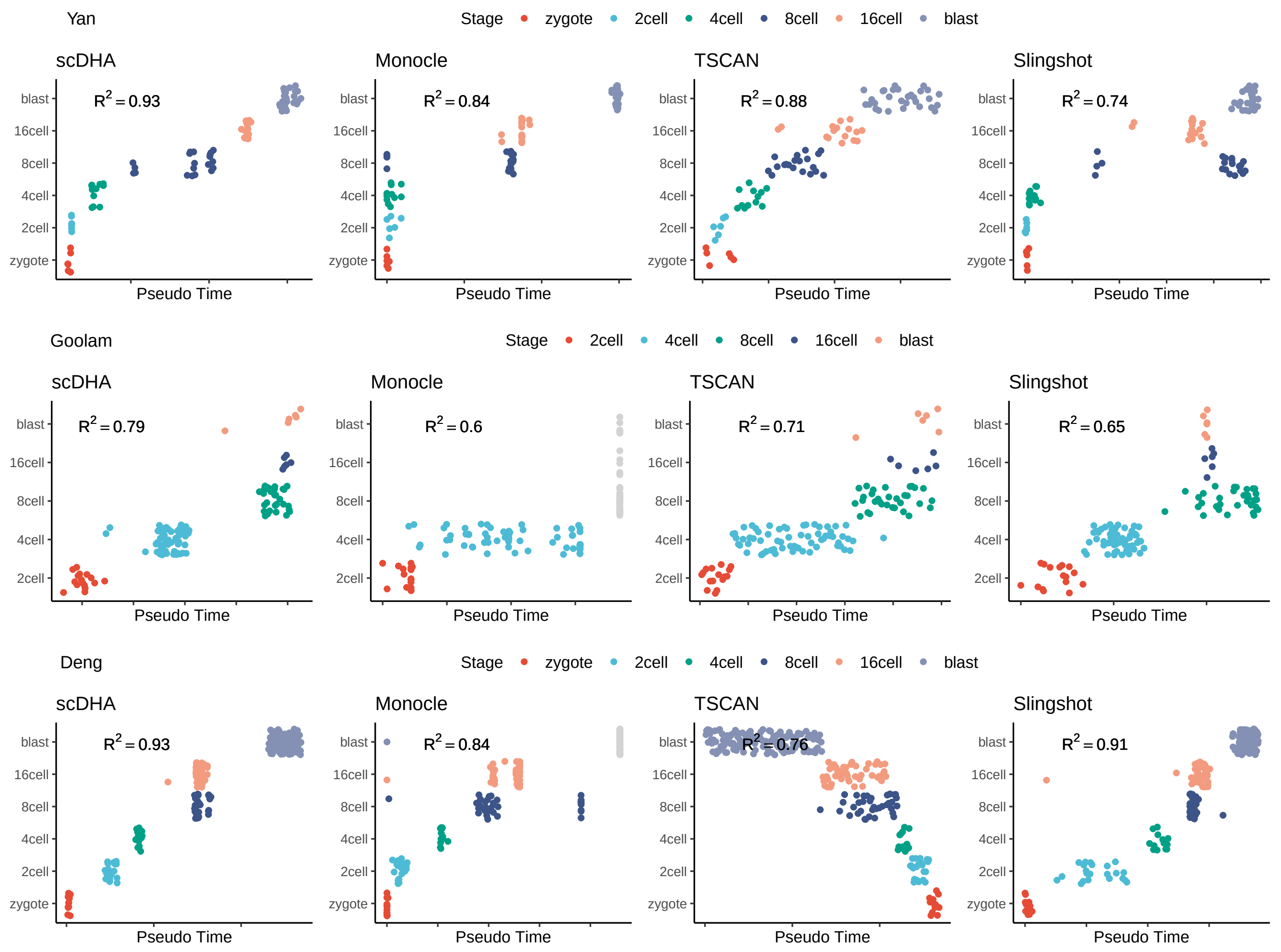


**Figure 8.** Pseudo time data inferred from Yan, Goolam, and Deng dataset using scDHA, Monocle, TSCAN and Slingshot. $R^2$ is coefficient of determination between inferred pseudo time and development stage. Points with gray color mean cells with infinity pseudo time from Monocle.

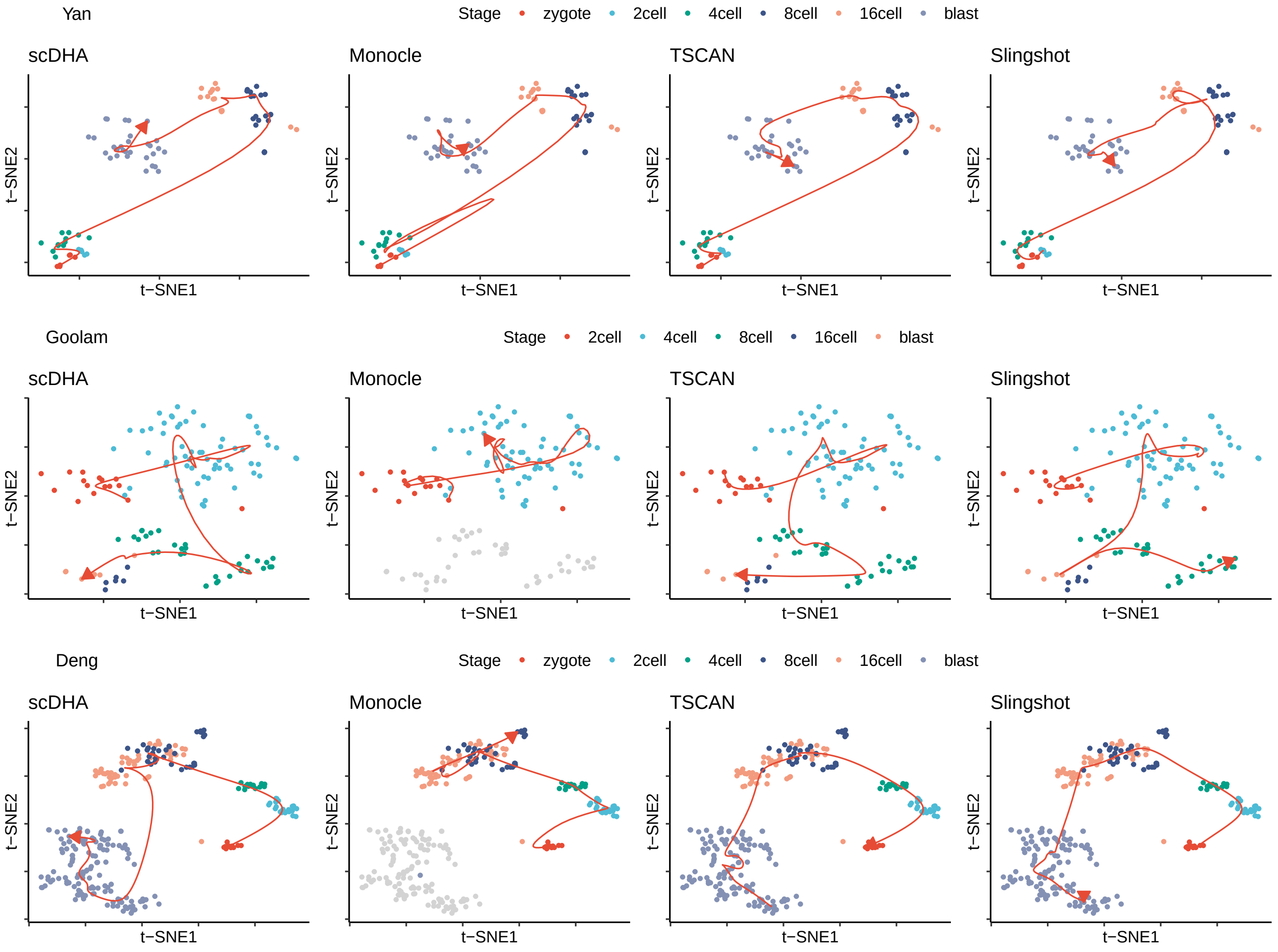


**Figure 9.** Visualized trajectory inferred from Yan, Goolam, and Deng dataset using scDHA, Monocle, TSCAN, and Slingshot. Points with gray color mean cells with infinity pseudo time from Monocle.
